# Convergent evolution of red pigmentation in extrafloral nectaries: global patterns and mechanisms

**DOI:** 10.64898/2026.02.24.707775

**Authors:** Sylvie Martin-Eberhardt, Vincent S. Pan, Kadeem J. Gilbert, Marjorie G. Weber

## Abstract

Convergent traits are a signature of adaptation, where shared selective pressures may lead to the replicated evolution of traits with similar function. Here, we present an integrative investigation of a widespread yet understudied feature of an iconic, convergent mutualistic plant trait – red pigmentation in extrafloral nectaries (EFNs). We surveyed 625 species to describe the distribution of red coloration in EFNs across plant morphology, biogeography, and phylogeny for the first time and propose several adaptive hypotheses for the drivers of their convergence across plants. We then test several of these hypotheses by integrating our global-scale biogeographic and phylogenetic analysis with phylogenetically paired common garden bioassays and a museum collections survey. We find that across the plant phylogeny, cold wet regions and deserts are hotspots of red EFN coloration, and multiple lines of evidence point to a fungal defense role of red pigments in EFNs. Ultimately, our findings highlight red EFNs as a highly convergent phenotype across plants and point to distinct selective pressures underlying the distribution of EFN coloration and EFNs themselves.

## Introduction

Convergent traits – those that have evolved multiple times independently across distantly related lineages – offer unique windows into the drivers and consequences of trait evolution. The evolutionary replication provided by convergent phenotypes creates a “natural experiment” to test hypotheses about the ecological and evolutionary consequences of a given trait, as well as the selective pressures that shape trait evolution (Powell and Mariscal 2015). One highly convergent, ecologically important trait in plants are extrafloral nectaries (EFNs), sugar secreting glands that attract arthropod predators to plant tissue whose presence and aggression reduce herbivory (Bentley 1977). EFNs are highly convergent across plants, arising in diverse plant tissues of at least 457 independent lineages across flowering plants (Weber and Keeler 2013, Marazzi 2013), and are highly studied as a paradigmatic plant defense trait and textbook example of mutualism. Here, we report on a striking yet largely uninvestigated feature of EFNs – their often conspicuously scarlet, red-orange, or maroon color (e.g. Figure 1A-D). EFNs with red pigmentation that contrasts with the color of surrounding tissue can be observed across plants. However, the phenomenon of red EFNs has been largely ignored, leaving the distribution and function of this striking pigmentation largely unknown.

**Figure 1.**
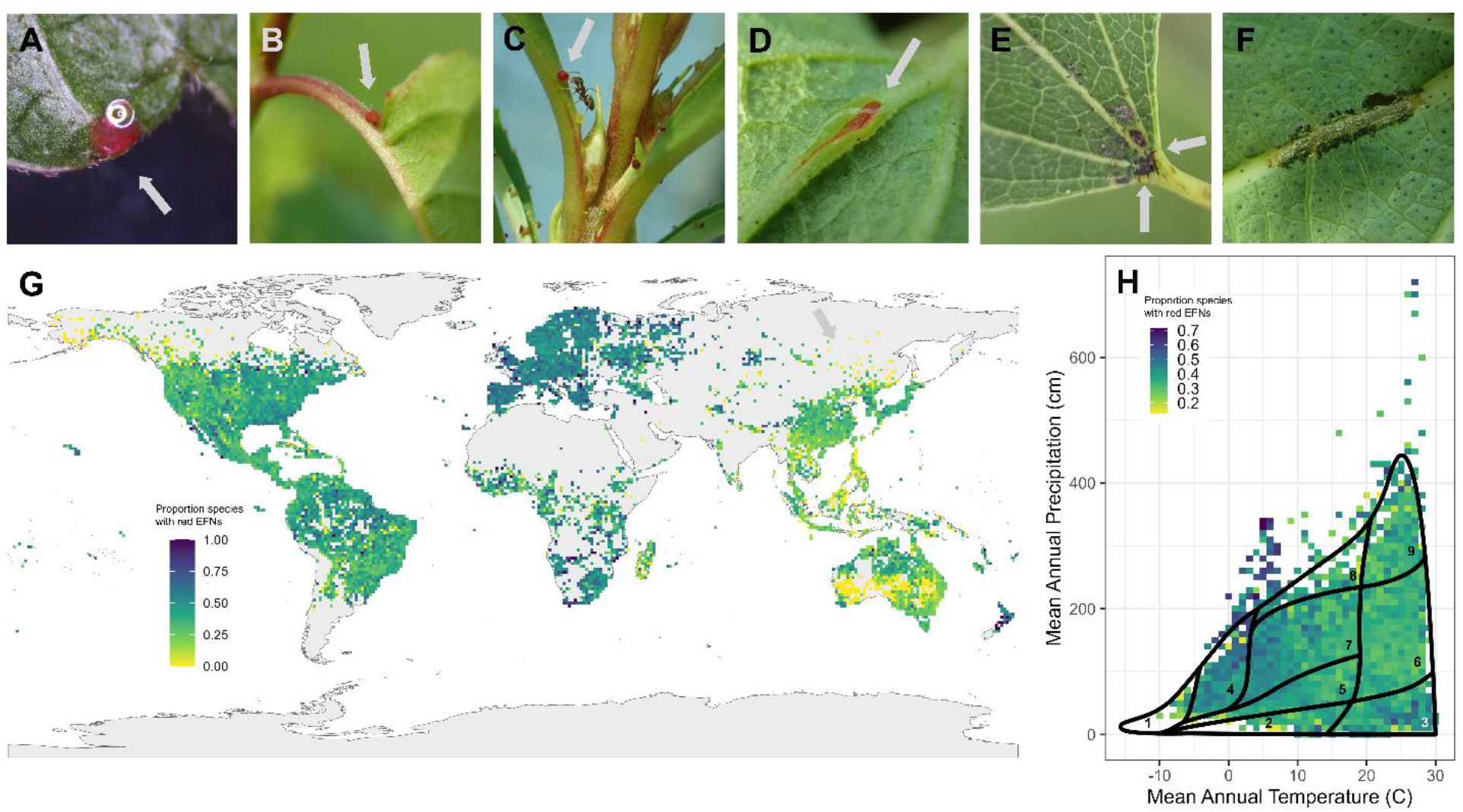
(A-D) Examples of red EFNs: A) *Paullinia pinnata* leaflet. B) *Prunus avium* leaf base. C) *Impatiens balsamina* petiole visited by ant. D) *Gossypium australe* midvein EFN. E) *Populus tremuloides* EFN infected with mold. F) Targeted herbivory on *G. sturtianum* EFN (by *Gryllodes sigillatus*). G) One degree raster map showing proportion of species with red EFNs in each raster containing three or more species (GBIF EFN color dataset). H) GBIF EFN color dataset plotted in biome space, with Whittaker biome plot overlaid. 1 = tundra, 2 = temperate grassland, 3 = desert, 4 = boreal forest, 5 = woodland/shrubland, 6 = tropical seasonal forest, 7 = temperate seasonal forest, 8 = temperate rainforest, 9 = tropical rainforest. Image credits: A) R. Glos, B-F) S. Martin-Eberhardt.

Here, we report on the biogeographic and phylogenetic distribution of red EFNs across plants, establishing the convergent nature of red coloration within EFN-bearing lineages using primary literature, iNaturalist, living plants, and field photos. We then integrate this dataset with biogeographic modeling, museum collection data, and experimental approaches to look for evidence supporting adaptive hypotheses underlying red EFNS convergence, touching on four of seven novel hypotheses for the pressures underlying their convergence, and (see Box 1, Table 1 – red EFN hypotheses). Together, our approach leverages the convergent nature of EFNs across plants to present and test potential hypotheses for the drivers of convergent pigmentation within an iconic convergent plant defense trait

**Table 1.** Adaptive hypotheses and resulting predictions for convergent red coloration of EFNs.

| Hypothesis | Description | Global Predictions | Ecological Predictions |
| --- | --- | --- | --- |
| Single selective pressure (herbivory) |  |  |  |
| Mutualist Advertisement | Red pigmentation makes EFNs easier to locate for mutualistic visitors | Red EFNs will be more common in areas of high visitation by red-sensitive mutualists (e.g. jumping spiders) | Red EFNs will be more quickly located or more frequently revisited by mutualists |
| EFN Warming | Red pigmentation absorbs warms nectary tissue and increases VOC emission | Plants in colder areas will be more likely to have red EFNs | Red EFNs will be warmer than green EFNs and emit a stronger scent under cold conditions, facilitating mutualist detection |
| Multiple selective pressures |  |  |  |
| Anti-fungal | Anthocyanins and betalains protect EFNs from fungal attack | Red EFNs will be more common in habitats with high fungal pressure | Red EFNs will experience reduced fungal infection |
| Anti-herbivory | Red pigmentation warns herbivores that EFNs are well-defended | Red EFNs will be more common in areas with high herbivore pressure | Red EFNs will experience reduced herbivory |
| Osmotic Stress | Anthocyanins and betalains mitigate osmotic stress from contact with sugary nectar | Plants that produce more concentrated nectars will be more likely to have red EFNs | Red EFNs will secrete nectar longer than green EFNs under high osmotic stress conditions |
| Photodamage | Anthocyanins and betalains protect EFN tissues from photodamage | Plants in areas with higher irradiance will be more likely to have red EFNs | Red EFNs will remain active longer than green EFNs under high irradiance |
| Either one or multiple selective pressures |  |  |  |
| Compound specificity | Different anthocyanins or betalains have different functions | Some pigment classes will be more common in biogeographic areas dominated by particular stressors | Different stressors will upregulate different groups of anthocyanins or betalains |

### Box 1: Adaptive hypotheses for red EFNs in plants

Phenotypic convergence in nature points to the potential for shared selective pressures underlying repeated evolutionary patterns. Here, we propose seven hypotheses for the adaptive value of red EFNs in plants underlying their convergence (Table 1). In plants, red and purple pigmentation is primarily caused by anthocyanins, diverse groups of plant pigments that vary in their hydroxylation and sugar decorations (Grotewold 2006). Betalains replace anthocyanins as red pigments within some lineages in the Caryophyllales and appear to serve parallel functions (Jain and Gould 2015*a*). Our hypotheses integrate biochemical, physiological, and ecological function of red pigmentation in plants with our understanding of EFNs as key traits in mutualistic interactions. The hypotheses fall into two broad categories: 1) those in which a single selective pressure (herbivory pressure) selects for both EFNs and their red pigmentation, and 2) those in which a distinct selective pressure arises in response to the emergence of the EFN trait (Box 1).

### Hypotheses proposing a single selective pressure

#### Mutualist Advertisement Hypothesis

We propose that red pigments in EFNs could serve a signaling role in interactions with mutualists (e.g. ants, spiders, and wasps (Cuautle and Rico-Gray 2003; de Oliveira Dias and Stefani 2024), similar to the role of red pigments in other plant parts such as fruit and flowers (Miller et al. 2011; Sinnott-Armstrong et al. 2021; Jaeger et al. 2023). In this scenario, red coloration is part of a larger EFN trait syndrome that maximizes the effectiveness of extrafloral nectar secretion by increasing mutualist visitation. While jumping spiders are the only group of EFN visitors known to have long wavelength receptors equipped for discrimination of red coloration (Morehouse 2020), mutualists without long wavelength receptors such as ants (Yilmaz and Spaethe 2022) and wasps (van der Kooi et al. 2021) may still perceive red as dark in contrast to surrounding green leaf or stem tissue resulting in a salient pattern (Cammaerts 2012). In addition, anthocyanins and betalains can reflect in both the red (long wavelengths) and blue (short wavelengths), and thus purple EFN pigmentation could serve as a chromatic signal for wasp and ant visitors. Support for this hypothesis would include faster detection or more frequent visitation of red EFNs by mutualists.

#### EFN Warming Hypothesis

Concentrated red pigments in EFNs could increase EFN temperature, volatilizing scent compounds and increasing their attractiveness to mutualists. Dark red pigmentation is proposed to absorb solar energy and warm plant leaves (Hughes 2011). Thus, warming in the EFNs could advertise nectar to mutualists if warming increases volatility of nectar VOCs, which are known to attract mutualists to both EFNs and floral nectars (González-Teuber and Heil 2009). If the EFN warming hypothesis is supported, we expect red EFNs to be concentrated in cold habitats, where a small increase in leaf surface temperature could lead to substantial increase in volatilization and ultimately, mutualist visitation.

### Hypotheses proposing multiple selective pressures

#### Anti-fungal Hypothesis

Because both anthocyanins and betalains have anti-fungal properties (Jain and Gould 2015*b*; Naing and Kim 2021), we hypothesize red pigments may protect EFNs from fungi exploiting their sugar resources, leading to the convergent evolution of red EFNs in many species. As antioxidants, anthocyanins defend against fungal attack by quenching infection-induced bursts of reactive oxygen species that can trigger cell death, thus slowing the spread of necrotrophic fungus that feed on dead cells (Barna et al. 2012). In addition, anthocyanins are also potent antifungals when deglycosylated, which occurs when fungal infection causes cell lysis (Sudheeran et al. 2020). Further, because EFNs concentrate sugars, EFN tissue may be particularly susceptible to fungal attack. Indeed, EFNs commonly become coated in sooty mold (Escalante-Pérez et al. 2012) and other sugar-feeding fungi (Figure 1E), which could impact their functionality in recruiting defensive mutualists. Finally, some EFNs are vascularized (Contreras and Lersten 1984; Rodríguez-Morales et al. 2016; Ávila-Argáez et al. 2019) providing a conduit into the plant for fungal pathogens. We predict that if anthocyanins and betalains serve an antifungal role in EFNs, they will be more common in habitats with greater fungal pressure.

#### Anti-herbivory Hypothesis

We propose that red coloration in EFNs could act as an anti-herbivory warning signal. In leaves, red coloration has been demonstrated to play an herbivore warning function, indicating well-defended tissues (Cooney et al. 2012; Portillo-Nava et al. 2021; Cornelissen et al. 2025). As a plant defense structure, EFNs are particularly valuable tissues for plants, yet their high sugar content also makes them particularly nutritious for herbivores to consume (e.g. Figure 1F).

Together, these dynamics may select for increased defenses in EFN relative to non-EFN tissue. Indeed, Gish *et al*. found higher levels of chemical and physical defense in fava bean EFNs compared to surrounding tissues (2015, 2016). Support for this hypothesis includes red EFNs experiencing reduced herbivory and occurring in habitats with greater herbivory pressure.

#### Osmotic Stress Hypothesis

Because concentrated solute solutions like nectar can cause osmotic stress on the leaf surface, we hypothesize that anthocyanins and betalains could mitigate osmotic stress in convergent, nectar-secreting EFN tissue. Sugar can induce osmotic stress (Cui et al. 2010), and sugar concentrations in extrafloral nectar are typically high (comparable to or higher than floral nectar) (Wunnachit et al. 1992; Koptur 1994). Thus, the cells at the surface of the EFN likely experience substantial osmotic stress from their own secretions. The reactive oxygen species generated under osmotic stress conditions can be quenched by anthocyanins and betalains, protecting the pigmented tissue (Jain and Gould 2015*a*). Under this hypothesis, we predict that EFNs that produce more concentrated nectars will be more likely to produce high concentrations of red, antioxidant anthocyanin and betalain pigments.

#### Photoprotection Hypothesis

We hypothesize that red pigments may prevent photodamage from solar radiation. In other plant parts, anthocyanins and betalains are upregulated by UV, have been demonstrated to reduce photo-oxidative damage (Jain and Gould 2015*a*), and are hypothesized to both attenuate non-photosynthetically active wavelengths and quench reactive oxygen species generated under light stress conditions (Nakashima et al. 2011; Landi et al. 2015). Particularly if chlorophyll concentrations are lower in EFNs, which is thus far unknown, EFN tissues could be uniquely susceptible to photodamage, making high concentrations of anthocyanins or betalains adaptive. Support for this hypothesis includes a greater prevalence of red EFNs in deserts and other regions with high irradiance.

### Either one or multiple selective pressures

#### Compound Specificity Hypothesis

Integrating the previous hypotheses, different anthocyanin or betalain compounds may serve different ecological or physiological functions for the plant according to their biochemical properties. An emerging hypothesis to explain why plants synthesize hundreds of unique anthocyanin compounds is that different types of anthocyanins perform different functions for the plant. Anthocyanins and betalains vary in their antioxidant activity (Belhadj Slimen et al. 2017; Dudek et al. 2022) and different stressors have been demonstrated to upregulate different specific anthocyanin compounds (Kovinich et al. 2014). As such, different selective pressures may favor different assemblages of anthocyanins in EFNs; for example, osmotic stress might favor highly hydroxylated anthocyanins with high antioxidant capacity, while photoprotection may favor a diversity of hydroxylation states that together reflect – and thus protect EFNs from – a wide range of wavelengths. More broadly, convergent expression of particular classes of anthocyanins or betalains in habitats dominated by particular stressors would support the compound specificity hypothesis.

Finally, red coloration in EFNs may not be adaptive at all, and some or all pigments in EFN tissue could be the result of pleiotropic or other genetic constraints. While the striking level of convergence in EFN redness in tissue with the same function from distantly related plants suggests an adaptive role, functional hypotheses should be tested, rather than an adaptive function assumed.

## Methods

We first investigated the phylogenetic and morphological distribution of red EFNs across plants using a combination of primary literature, inspection of living plants, and expert-identified images. We then accessed global herbarium records of species with known EFN color to identify biogeographic patterns of EFN coloration. Finally, we integrated several approaches to test for evidence consistent with four of the seven hypotheses presented above: the anti-fungal, anti-herbivory, photodamage, and EFN warming hypotheses. Using the global herbarium record dataset, we tested biogeographical hypotheses concerning how species with red or non-red EFNs are distributed across gradients of phytopathogen relative abundance, temperature, and irradiance; and creating a phylogenetically paired common garden for experimental comparisons of EFN fungal infection and herbivory rates.

### EFN color dataset: EFN color and morphology survey

To compile a dataset of EFN coloration we broadly surveyed EFN color across EFN-bearing plants using a combination of primary literature, image, and live plant sources. We targeted species known to possess EFNs as listed in the World List of Plants with Extrafloral Nectaries (Weber et al. 2023). Data on coloration was compiled from color photographs and written descriptions of nectary color in primary literature cited by the World List of Plants with Extrafloral Nectaries, images of fresh tissue published by Arboretum Wespelaar (De Langhe 2012), unpublished images collected and identified to species by clade experts (Phyllis Coley, pers. com., Pedro A.P. Rodrigues, pers. com., Pooja Nathan, pers. com.), and research-grade images from iNaturalist. For iNaturalist images, we scored three to five images per species and counted the species as red if at least one image contained red EFNs, indicating the species could create EFNs with concentrated anthocyanins or betalains. We also scored EFN color from live plants in botanic gardens, university collections, or from natural populations (identified by [author name removed for double-blind review], see supplemental data); EFN color records from living plants were made only from fresh nectaries.

To efficiently categorize EFN color from a range of sources, two observers scored EFNs as green, yellow, orange, scarlet, maroon, brown, or black; each species could be added to more than one color category to reflect variation in EFN color. As a conservative categorization of EFNs with a high concentration of anthocyanins or betalains, “red EFN plants” were those with records of scarlet or maroon nectaries; “green EFN plants” were all others. While carotenoids can contribute to red coloration (Grotewold 2006), they also function as antioxidants and even antifungals (Orsi et al. 2023); thus inclusion of any red EFNs pigmented primarily by carotenoids in the dataset would not confound our hypotheses. To check for differences in color scoring calls, the rate of red EFN detection was compared for the two observers.

To detect differences in EFN coloration in different parts of the plant, we also recorded EFN location, which was binned into one of eight categories (per Figure 2B). Plants with nectaries in distinct locations could be assigned to more than one location category, each with a separate EFN color entry. See supplement for category details.

**Figure 2.**
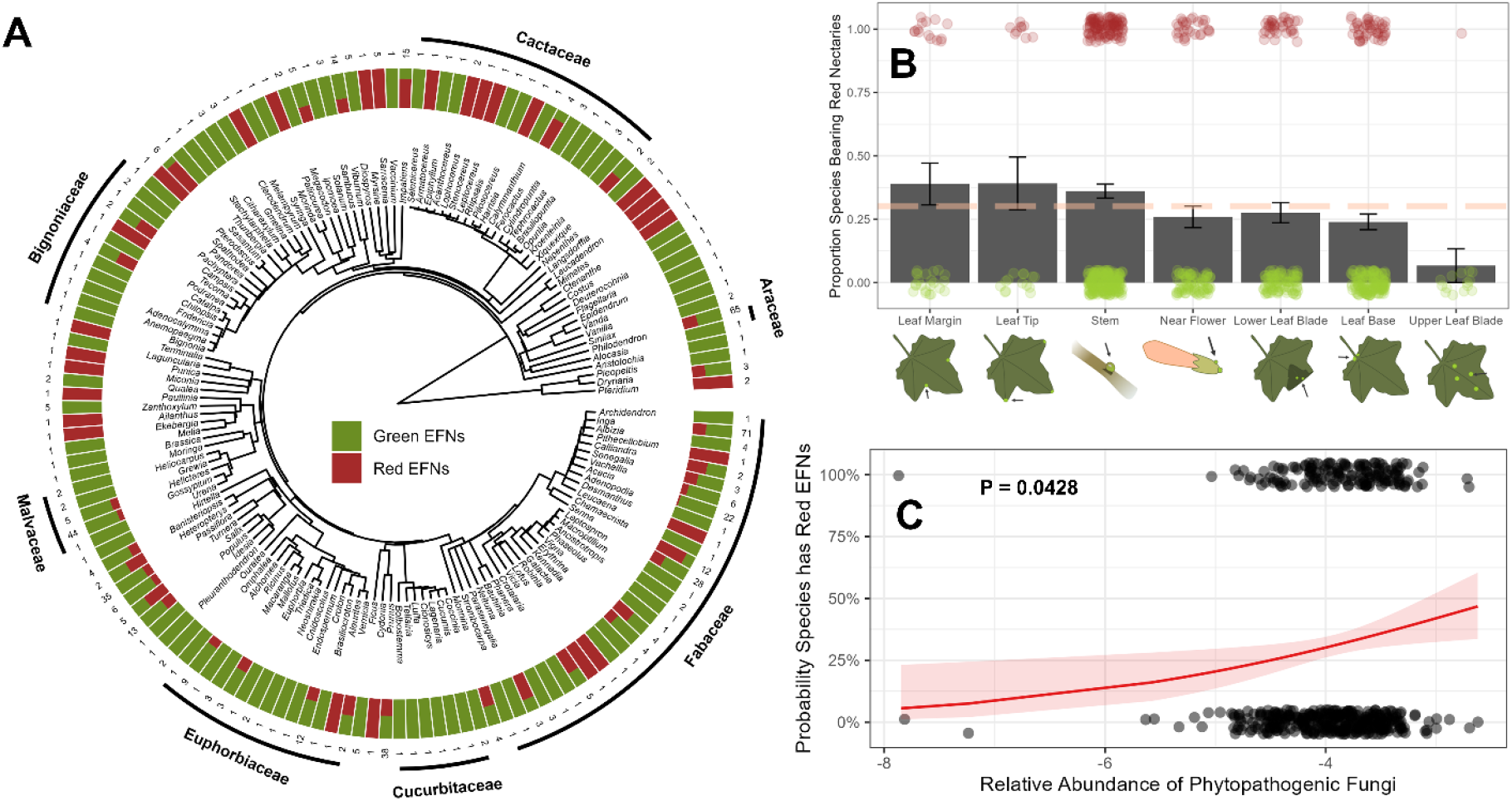
A) Phylogenetic tree showing all sampled genera with proportion of species with red EFNs and total species sampled in each genus plotted at tips. B) Nectaries on the leaf margin, leaf tip, and stem had a higher rate of red pigmentation than the overall mean rate (dashed line), while those on the upper leaf blade were below the mean rate. C) Predicted relative abundance of fungal phytopathogens across a species’ range predicts a marginally significant increase in probability a species will have red EFNs.

### GBIF Dataset

We examined the global pattern of red EFN occurrence by downloading all herbarium records of each species in our EFN color survey from the Global Biodiversity Information Facility (GBIF) using *rgbif* (Chamberlain et al. 2017). To harmonize taxonomic names, we used the Online Matching Tool from World Flora Online (Kindt 2025) with 3-character fuzzy matching; for taxa with no valid name, we defaulted to the authority used in the paper, and then to the oldest valid record. This yielded a 97.9% matching success rate. We excluded all records not at the species level and all cultivated records. To clean the georeferenced coordinates for use in establishing climatic conditions across the species range, we excluded all records in water bodies, Antarctica, country capitals, centroids, institutions, and urban areas, as well as records with equal latitude and longitude and those with impossible coordinates using the *CoordinateCleaner* package (Zizka et al. 2019). Finally, we used the median absolute distance method implemented in *CoordinateCleaner* to remove implausible outlier records.

To project the distribution of red EFNs into biome space, we extracted mean annual temperature (MAT) and precipitation (MAP) for each specimen (see EFN warming and photodamage hypothesis section for details). We rounded all records to the nearest 1°C mean annual temperature and 10 cm mean annual precipitation to create raster cells in biome space. After thinning to one record per species per raster cell, we then plotted raster cells with five or more species onto Whittaker biome space colored by proportion of species with red EFNs.

### Herbarium Dataset

To test for an effect of red coloration on fungal infection of EFNs, we examined herbarium specimens for fungal hyphae on EFNs (see Anti-fungal Hypothesis methods section below for details). We selected pairs of red- and green-EFN-bearing species in the same genus, prioritizing pairs from the same geographic region and with the same EFN morphology. We inspected 7-15 specimens per species from each of 15 congener pairs representing 11 families, (435 specimens total, Table S1).

### Common Garden Bioassays

We experimentally tested for a difference in susceptibility to fungal infection or herbivory using a common garden composed of three of the congener pairs used in the herbarium study: *Passiflora, Sesamum*, and *Viburnum* (see Table S1). Exceptions included the use of *S. trilobum* (instead of *S. triphyllum*) based on horticultural availability. We also included an intraspecific comparison using *Impatiens balsamina*, which displays distinctive within-species variation in EFN color. Plants were grown in 15% sand, and started in greenhouse or growth chamber conditions and moved to an outdoor, fenced common garden in Michigan, USA (42.410, - 85.391) in May and June 2026. To accommodate the growth conditions required for different species, *Viburnum* and *Sesamum* plants were grown in full sun, while *Passiflora* and *Impatiens* were grown under 42% shade cloth. With the exception of *S. trilobum* (N = 13), 24-30 plants per taxon were used for all bioassays.

### Common Garden Anthocyanin and Chlorophyll Quantification

Because chlorophyll can mask anthocyanins and other pigments, it is unknown whether red EFNs have a higher concentration of red pigments than surrounding tissue, or whether they simply lack chlorophyll compared to nearby tissue and thus appear redder. Thus, we quantified both chlorophyll and anthocyanin concentrations in EFNs to test if scarlet and maroon tissues had higher concentrations of anthocyanin than green tissues. We compared the anthocyanin concentrations in red EFN tissues compared to neighboring tissue in two species: *Viburnum dentatum* and *I. balsamina* (red EFN variety). We also compared red and green EFN tissue across common garden species. For all extractions, we harvested EFNs or adjacent tissue, pooling tissue to achieve sufficient biomass, then weighed pooled tissue, and immediately ground it in acidified methanol (1% HCl, 80% Methanol, 19% RO H_2_O). To eliminate turbidity, samples were centrifuged for 10 minutes at 4000 rpm, then held at 4°C. 300 uL of each extract was added to a flat-bottom 96-well plate, which was then read at 535 nm in a BioTek Synergy H1 plate reader (Agilent, Santa Clara, USA). Anthocyanin concentration was calculated from a standard curve of cyanidin-3-glucoside (Sigma Aldrich, St. Louis, USA). To compare the relative levels of chlorophyll in EFNs across our common garden species, we harvested EFNs using the same procedure and pool size and ground harvested tissue in 96% ethanol, then incubated samples overnight at 4°C before centrifugation and spectrophotometry at 645 nm. Chlorophyll absorbances were then standardized by tissue mass.

## Hypothesis Testing

### Anti-fungal hypothesis

To test the anti-fungal hypothesis, we combined the global herbarium record dataset (extracted from GBIF) with the phylogenetically paired survey of infection in herbarium specimens and the phylogenetically paired bioassay in common garden plants.

#### Global data

We tested for a broadscale geographic association between fungal phytopathogen pressure and red EFN prevalence. To estimate fungal phytopathogen pressure at a global scale, we trained a gradient boosting machine (GBM) on 13,911 records of global ITS sequencing data with associated phytopathogen hypotheses (Li et al. 2023). The model predicted the relative abundance of fungal phytopathogens with an out-of-sample predictive performance of r^2^ = 0.51 (mean absolute error = 0.060 proportion phytopathogen), indicating a reasonable ability to predict site level fungal phytopathogen risk. See supplement for details.

To obtain an estimate of phytopathogen pressure for each record in our GBIF dataset (occurrence records of plants scored in the EFN color survey), we predicted fungal pathogen relative abundance from the trained GBM, excluding observations too dissimilar to the training dataset (dissimilarity index < 0.4, Meyer and Pebesma 2021). We then calculated the median relative phytopathogen abundance experienced across the range of each species (thinned to one record per species per 2.5 minutes latitude and longitude) with five or more records to summarize the phytopathogen pressure experienced by each species. We included fungal pressure, mean annual temperature (MAT), and irradiance as predictors of EFN color, allowing us to compare the relative importance of fungal pressure with the EFN warming and photodamage hypotheses (see below).

#### Analysis of herbarium dataset

We tested for an effect of red coloration on fungal infection of EFNs by inspecting herbarium specimens in our phylogenetically, geographically, and morphologically paired herbarium dataset for fungal hyphae on EFNs. To minimize incidence of mold growth after collection, we used the youngest available specimens and scored only black hyphal growth consistent with sooty mold, which grows on EF nectar (Torres-Hernández et al. 2000) and can reduce photosynthesis (Lemos Filho and Paiva 2006; Santos et al. 2013). On each specimen, we used a dissecting scope to score the number of infected or consumed nectaries out of the total number present at up to 10 locations. We also extracted the fungal phytopathogen relative abundance prediction for each georeferenced herbarium record to use in addition to binary EFN color as predictors for the rate of EFN infection.

#### Common garden fungus bioassays

To experimentally test for an effect of anthocyanin pigments on rates of EFN infection, we inoculated EFNs in our common garden with grey mold (*Botrytis cinerea*), a globally distributed extreme generalist necrotrophic fungus (Chen et al. 2023), and asked whether EFN color predicted resulting fungal growth. We applied *B. cinerea* isolated from Michigan commercial grape production. To simulate herbivory or other incidental damage, we pierced each nectary on 3-10 leaves per plant (13-30 plants per taxon × eight taxa = 1013 leaves total) that were intact and not senescing with a fine sewing needle before using a paintbrush to apply sporulating *B. cinerea* to the EFN. Needles were cleaned with ethanol between each plant. After a 12-day growth period, inoculated leaves were harvested for inspection under the dissecting scope at 40x magnification for fungal hyphae.

#### Anti-herbivory hypothesis

We tested the herbivore warning hypothesis by combining herbarium specimen surveys with the insect bioassay. We surveyed 435 specimens from 32 species and 11 families (herbarium infection dataset) for targeted herbivory of EFNs, identifying chewed and missing EFNs using a dissecting scope; within each pair. To create an herbivory bioassay, we harvested one whole leaf (or inflorescence, in the case of *Sesamum*) from each common garden plant (described under the Anti-fungal Hypothesis). These leaves were paired in a humidified petri dish (14 cm diameter) with a leaf or inflorescence of their congener or intraspecific pair. To maximize extrafloral nectary production and availability, leaves were allowed to humidify for five hours before the start of the experiment, and the number of EFNs was recorded for each leaf. A single tropical house cricket (*Gryllodes sigillatus*) that had been starved for six and a half hours was placed in each petri dish. After six days of feeding, we examined each nectary for damage and incidental fungal infection under a dissecting scope. Crickets were chosen as a model herbivore based on their status as generalists and findings of Gish *et al*. (2015) that crickets selectively feed on EFN tissue.

#### EFN warming and photodamage hypotheses

To test the EFN warming and photodamage hypotheses, we tested for a relationship between the probability of having red EFNs and the median irradiance and temperature across each species’ range. We calculated the median MAT and irradiance for each species in our GBIF dataset, using the same approach as the fungal phytopathogen data above. In brief, we accessed bioclimatic variables (WorldClim, Fick and Hijmans 2017) at 10 minute resolution using the *geodata* package (Hijmans et al. 2024), and extracted the MAT and irradiance for each herbarium record (thinned to one record per species per 10 minutes latitude and longitude) using the *terra* package (Hijmans et al. 2022).

### Statistical methods

All analyses were performed in R version 4.5.0; we used the package *phyr* for all analyses with a phylogenetic random effect, except for the biogeography analysis of EFN color, which was run in a Bayesian framework with *brms* (Bürkner 2017). All phylogenetic trees for models with phylogenetic random effects were built using the *V*.*phylomaker* package. Analyses without phylogenetic random effects were fit with glmmTMB (Brooks et al. 2017).

Throughout, binary EFN color and EFN infection (number infected out of total) were modeled with a binomial distribution. Phylogenetic relatedness along with plant (common garden data) or specimen (herbarium data) identity were used as random effects. To model fungal infection and herbivory in our common garden bioassays, we omitted phylogenetic random effects because of the small number (n = 4) of distantly related genera each in its own order (Ericales, Malpighiales, Lamiales, and Dipsacales) limits the possibility of substantial phylogenetic non-independence effects. To model cricket bioassays (herbivory and incidental EFN infection), we used EFN color as a predictor and arena as a random effect nested within genus.

## Results

### EFN color and morphology survey

In total, we scored 625 species from 154 genera and 54 families, finding 31.0% of all species scored could produce red EFNs. The two observers recorded similar rates of red EFN scores: 28.8% vs 30.8% with red EFNs. Red EFN producing species were phylogenetically widespread (Figure 2A), with 56.6% of all families surveyed containing at least one red EFN producing species. We detected weak phylogenetic signal (*D* = 0.816, Fritz and Purvis 2010) indicating that more closely related species are slightly more likely both have red EFNs than expected by chance. Red EFNs occurred on all scored parts of the plant (Figure 2B); rates of red EFNs were lower on the upper leaf blade compared to leaf margin (P = 0.0508), leaf tip (P = 0.0546), and stem (P = 0.0549), although these differences were only marginally significant.

### Biogeography

Species producing red EFNs were widely distributed across the globe (Figure 1G). Species with red EFNs were more common than species with green EFNs in several geographic zones, including eastern North America, Amazonia, southern Africa, western and central Eurasia, and New Zealand. Conversely, red EFNs were absent in our dataset across the northern tundra as well as parts of the Australian desert and tropical Oceania. Projecting EFN distributions into biome space revealed that species bearing red EFNs clustered in two distinct biomes: 1) desert and adjacent hot, dry regions, and 2) cold, wet regions (of the boreal forest, temperate rainforest, and temperate deciduous forest) (Figure 1H). MAT and MAP interacted to predict proportion of species with red EFNs. The odds of having red EFNs was reduced by 12.2% (95% Credible Interval: [9.51%, 13.9%]) for every standard deviation increase in MAT, although this effect was dependent on the MAP; a standard deviation increase in MAP predicted a 7.69% reduction (95% CI: [3.92%, 10.4%]) in the effect of MAT on the odds having red EFNs. At the average of MAT, every standard deviation increase in MAP increased the odds of having red EFNs by 4.08% (95% CI: [1.01%, 7.25%]). This negative interaction is consistent with red EFN species hotspots in both hot, dry and cold, wet regions.

### Common Garden Anthocyanin Quantification

Comparing between EFN tissue and adjacent tissue from plants in our common garden experiment, visually red EFNs had an average anthocyanin concentration 4.57 times that of surrounding tissue (Figure 3A, P < 0.0001) and a slightly higher concentration of chlorophyll (Figure 3B, 1.23 times greater in EFNs than surrounding tissue, P = 0.294) in both *I. balsamina* and *V. dentatum*. This result indicates that conspicuously red EFNs can arise from a higher concentration of anthocyanin rather than from reduced chlorophyll concentration in EFNs that reveals anthocyanins otherwise masked in surrounding tissues.

**Figure 3.**
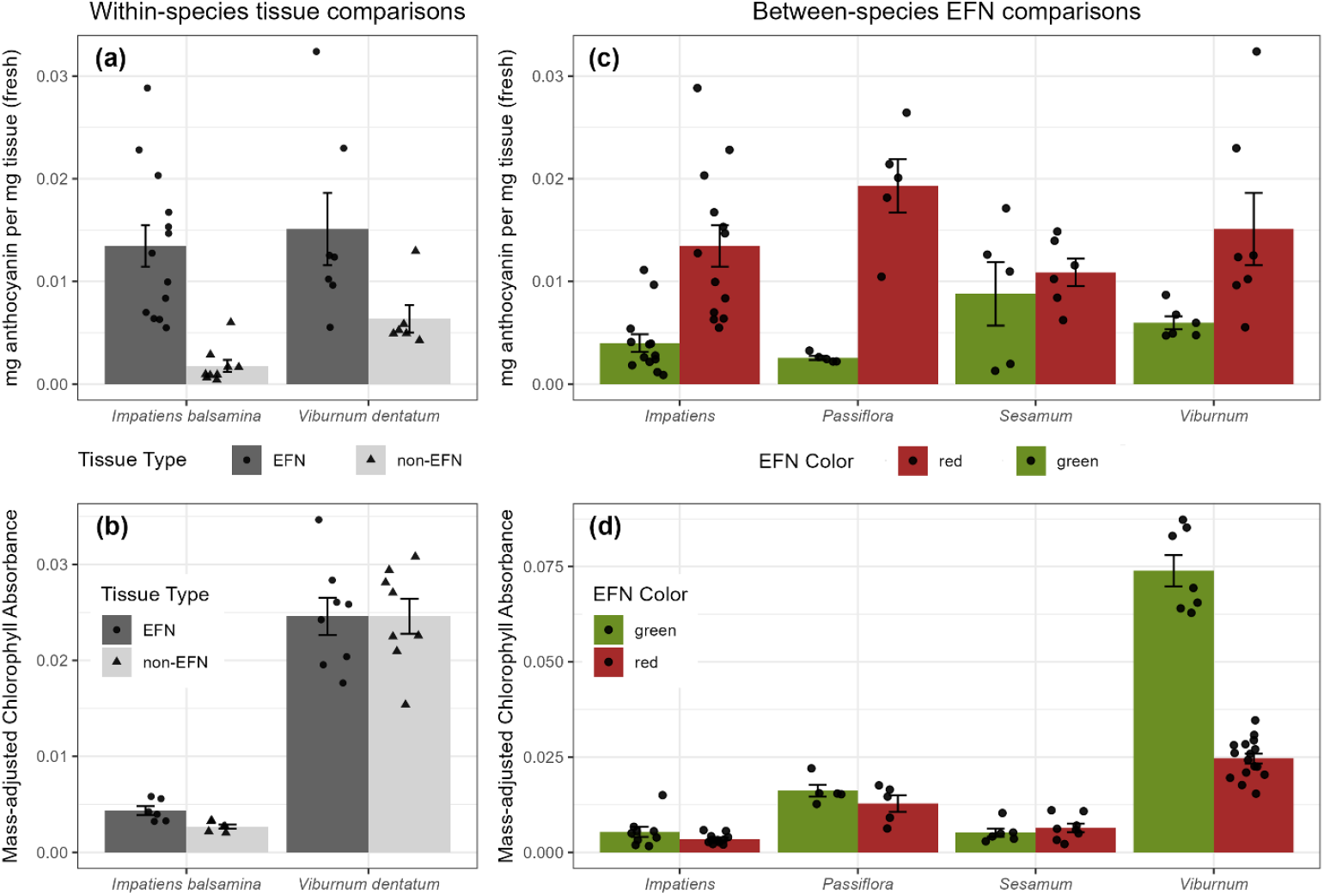
Biochemical analysis of EFNs and surrounding tissue. Comparing between EFN and neighboring tissue, EFNs had A) higher concentrations of anthocyanin and B) similar concentrations of chlorophyll compared to neighboring tissue for both *Impatiens balsamina* (red EFN variety) and *Viburnum dentatum*. C) In all species pairs surveyed, species with conspicuously scarlet or purple EFNs had higher concentrations of anthocyanins in those tissues, while D) levels of chlorophyll were largely equal within congener pairs.

Comparing EFN anthocyanin concentration in red- and green-EFN species within congener pairs, we confirmed that species chosen for their conspicuously red EFNs had higher levels of EFN anthocyanin compared to congeners with visibly green EFNs (Figure 3C, P < 0.0001). In addition, the concentration of chlorophyll in EFNs was typically not different between red- and green-EFN congeners (Figure 3D), indicating that red coloration was due to increased synthesis of red pigment rather than masking of red pigments by chlorophyll. The exception to this trend was the *Viburnum* congener pair, in which the chlorophyll concentration of *V. rafinesqueanum* (green EFNs) was three times greater than that of *V. dentatum* (red EFNs).

However, there was no overall effect of EFN color on chlorophyll concentration (P = 0.156) and the difference between *Viburnum* species is likely attributable to the fact that we compared leaves at different developmental stages due to horticultural availability of these slow-growing perennials; leaves of *V. rafinesqueanum* (green EFNs) were older than leaves of *V. dentatum* (red EFNs).

### Hypothesis testing

#### Anti-fungal hypothesis

In our global dataset of EFN coloration and species distribution, we found some support for the anti-fungal hypothesis. Every standard of deviation increase in median fungal phytopathogen relative abundance across a species’ range increased the odds of having red EFNs by 1.25 times (P = 0.0428). The relationship between fungal pressure and EFN coloration was stronger without the phylogenetic random effect (Figure 2C, β = 1.65, P = 0.0093), implying certain clades may be driving this relationship. Indeed, faceting by family revealed strong positive relationships in the well-sampled Bignoniaceae, Fabaceae, Salicaeae, Euphorbiaceae, and Malvaceae, while Passifloraceae, Rosaceae, and Araceae had a negative relationship between fungal pressure and EFN color (Figure S2).

In our herbarium infection dataset, EFN color did not predict infection (P = 0.420, Figure 4A), nor did it interact with phytopathogen relative abundance across the species range (P = 0.934).

**Figure 4.**
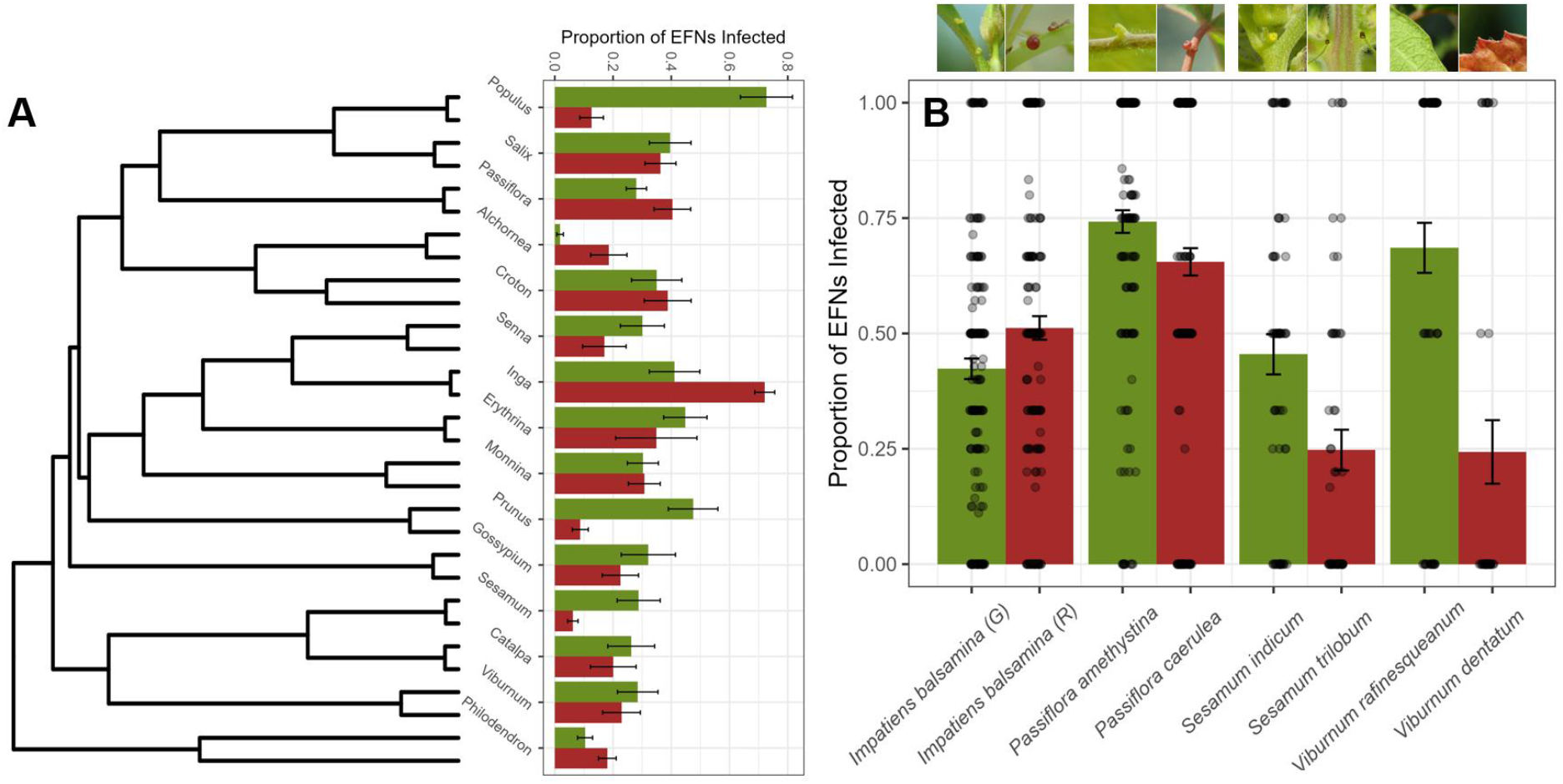
A) Fungal infection in herbarium specimens was not overall higher in species with green EFNs compared to congeners with red EFNs. B) Across four congener species pairs with red- and green-EFN species, green EFNs were infected at a higher rate than red EFNs for three of the four paired taxa.

In our common garden experiment, three of the four species pairs showed reduced infection in the red EFN species, although the overall effect of EFN color on infection was not significant (Figure 4B, P = 0.156). However, in the cricket bioassay, green EFNs were 1.42 times more likely to experience incidental infection compared to paired leaves with red EFNs (Figure S3, P = 0.030).

#### Anti-herbivory hypothesis

We found no support for the anti-herbivory hypothesis. Species with green EFNs were not more likely to be eaten by crickets than their red-EFN congener (P = 0.390). We also observed targeted herbivory of *Sesamum indicum* nectaries by a generalist Tortricid caterpillar (*Argyrotaenia velutinana*) in our common garden plants. In addition, within our herbarium dataset, we found just 13 instances of targeted nectary feeding; among these instances, 10 were in species with red EFNs, and the remaining 3 were in species with green EFNs.

#### EFN warming and Photodamage hypothesis

We found no support for the EFN warming hypothesis or the photodamage hypothesis. Neither median MAT nor mean annual irradiance across the species’ range predicted the odds that the species would have red EFNs (MAT: P = 0.996, log irradiance: P = 0.224).

## Discussion

We describe the biological distribution of widespread convergence in a plant defense trait – red pigmentation in extrafloral nectaries (EFNs) – and test for evidence in support of adaptive hypotheses underlying their evolution. We created a phylogenetically diverse dataset of EFN color spanning over 600 species and examined the phylogenetic, morphological, and biogeographic distribution of red EFNs for the first time. We found that red EFNs are widespread across plant phylogeny, morphology, and biogeography, with 30% of all species surveyed able to produce red EFNs. At the biochemical level, red coloration in EFNs appears to be driven by the concentration of red pigments rather than differences in chlorophyll concentrations that mask or reveal these red pigments. Two biome hotspots of red EFNs emerged: cold, wet regions and hot, dry regions. Our global analysis and bioassays point to antifungal defense as an adaptive function of red EFNs. More broadly, our results suggest that the drivers of EFN coloration are separate from the primary driver of EFN evolution itself.

We found several lines of evidence for red pigment serving a defensive role against fungi in EFNs. Red EFNs were predicted by fungal phytopathogen relative abundance globally, and in the common garden inoculation bioassays, three of the four species pairs suffered higher infection in the green EFN species. In addition, the herbivory bioassay provided further support for the anti-fungal hypothesis, with green EFNs suffering higher incidental infection. This bioassay mimicked natural paths of infection (wounding by insects) and occurred in a high humidity environment, likely facilitating an overall higher rate of infection establishment that could reveal patterns of fungal defense (Park 1990). While these experimental and biogeographic results point to an anti-fungal function of red EFNs, our herbarium infection dataset did not show phytopathogen pressure predicting EFN color. It is possible that red pigmentation is not effective against sooty mold growth on drying specimens, or that heavily infected branches were passed over for more pristine specimens during collection. Alternatively, the anti-fungal function may be pigment-specific (the compound specificity hypothesis), obscuring a general pattern across the herbarium specimens. Anthocyanins and betalains vary in their antioxidant function (Belhadj Slimen et al. 2017; Dudek et al. 2022); for example, anthocyanins with greater hydroxylation such as delphinidins and petunidins may be more effective at quenching reactive oxygen species. In addition, fungal identity, which was not quantifiable from herbarium specimens, may be a critical predictor of anthocyanin anti-fungal benefits. Antioxidants such as anthocyanins and betalains are primarily beneficial in defending against necrotrophic fungi, because the pigments quench the burst of reactive oxygen species triggered during infection, preventing the cell death that necrotrophs thrive on (Barna et al. 2012). Cell death is advantageous when defending against a biotrophic fungal pathogen feeding on living cells, but is detrimental against necrotrophs. We were unable to distinguish biotrophic and necrotrophic fungal infection on herbarium specimens, potentially obscuring the benefit of anti-fungal pigments in EFNs. Together, the concurrence of global biogeographic pattern and experimental data point to an anti-fungal role of red pigments in EFNs, perhaps contingent on pathogen type, specific climatic conditions, compound identity, or other factors.

The possibility of two distinct hotspots of red EFNs is somewhat surprising, but one explanation of this disjunct pattern may be that a combination of different selective pressures favor red-pigmented EFNs and vary across habitats. For example, hot spots in arid environments may be due to drought-induced osmotic stress being exacerbated by sugar-induced osmotic stress within the EFNs, especially if nectars become concentrated due to evaporation. In these desert environments, synthesizing high concentrations of antioxidant anthocyanin and betalain compounds could mitigate osmotic stress, drought stress, and fungal attack. In cold environments, osmotic stress within the EFN may compound with generalized cold stress, making accumulation of antioxidant pigments in the EFN particularly advantageous both for mitigating abiotic and fungal stressors. Although we were not able to test the osmotic stress hypothesis in this study, we suggest that osmotic stress may interact with other stressors, producing the observed pattern of two distinct hotspots of red EFNs in cold, wet regions and deserts.

While we found several lines of evidence in support of the anti-fungal hypothesis, the anti-herbivory, photodamage, and EFN warming hypotheses were not supported by our preliminary analyses. However, additional, fine-scale studies, especially of EFN VOC emission, temperature, or photosynthetic physiology may reveal subtler patterns obscured by our global biogeographic analysis. Going forward, incorporating data on EFN longevity, vascularization, and nectar removal rate could reveal clearer patterns of EFN biological defense and physiological stress. We also suggest tests of the mutualist advertisement hypothesis could leverage some of the dozens of species we recorded with ‘inverse contrast’ – red-pigmented tissue surrounding a green EFN. If the contrast between colors is increasing apparency to mutualists, species with ‘inverse contrast’ should also enjoy higher visitation rates than EFNs of any color that match the coloration of surrounding tissue. In addition, we found multiple examples of targeted herbivory of EFNs in herbarium specimens and suggest herbarium records as a fruitful avenue for exploring targeted herbivory of EFNs more generally. In contrast, our findings did not mirror the observations of Gish *et al*. that crickets target EFN tissues for consumption (2015), with targeted herbivory observed only in *Gossypium*, which was not used in the main herbivory bioassay due to poor germination. While EFN color does not appear to impact targeted herbivory of EFN tissues, we propose a general investigation into this uniquely disruptive plant-insect interaction.

Our results for red EFN color are markedly different from those of other large-scale analyses of red plant color: fruit coloration (Sinnott-Armstrong et al. 2021), fall leaf color (Renner and Zohner 2019), and flower color (Dellinger et al. 2024). In fruits and flowers, colder temperatures predicted more contrasting (darker) colors (Sinnott-Armstrong et al. 2021; Dellinger et al. 2024), while in EFNs, cold temperatures did not predict increased coloration.

Instead, dark (red) EFNs occurred in two distinct hotspots, one of which was the hottest biome on earth. In addition, patterns of global autumn leaf coloration were opposite those of EFN coloration, with Europe having the lowest share of species with red autumn leaves (Renner & Zohner 2019) but the highest proportion of red EFNs, compared to other northern regions. In addition, irradiance, which Renner & Zohner link to autumn leaf coloration (2019), did not predict EFN coloration. One notable similarity between flower color and EFN color is the dominance of purple and maroon coloration in sub-Saharan Africa (Figure S4, Dellinger et al. 2024 Figure 4b), although causes of this pattern for either flowers or EFNs remain enigmatic. More generally, EFN activity represents a longer phenological period than ephemeral flowers, fruits, or senescing leaves, consistent with differing patterns of global pigmentation. Overall, the differences between global fruit, autumn leaf, flower and EFN coloration patterns suggest separate selective pressures acting on these tissue types.

More broadly, our study suggests that red pigmentation in EFNs may have evolved in response to unique challenges plants face after evolving nectar secretion, a case of a primary phenotype resulting in novel costs that select for the evolution of a secondary phenotype. In other words, the evolution of EFNs – while benefitting plants by reducing herbivory – may have come with new challenges for plants, including increased osmotic pressure and susceptibility to fungal attack, for which red pigmentation is a (at least partial) solution. This stepwise evolutionary view of convergence, if true, has several implications for our understanding of EFN evolution and ecology. First, while EFNs are highly convergent – having evolved in hundreds of separate lineages (Weber and Keeler 2013) – their evolution may ultimately be constrained by lineages’ ability to evolve solutions to osmotic and fungal stressors, especially in environmental contexts where these stressors are severe. Second, a sequential viewpoint begs the question of whether the evolution of red pigmentation itself comes with any novel costs, and whether those costs lead to the evolution of additional levels of convergence in EFN phenotypes (does the development of the secondary trait expose the organism to a third selective pressure, and so on). For example, darkly pigmented EFNs may camouflage small beetle herbivores or the dark heads of caterpillars from predators (Koptur 1992), and thus defense from fungus might intensify herbivory. Finally, over two-thirds of the plants investigated had green, rather than red EFNs. Future work should investigate whether lineages without red pigmentation in their EFNs have invested more heavily in alternative strategies to mitigate osmotic and fungal stressors, such as antifungal compounds in their nectar (Escalante-Pérez et al. 2012), especially in high-stress environments. Ultimately, testing whether convergent features of EFNs show evidence of being sequential solutions (identifying a pattern of one trait altering selective pressures that drives the evolution of another trait) will inform on the evolution of both the primary (EFN) and secondary (red coloration) phenotypes.

## Conclusion

In sum, we established that red coloration is highly convergent within an ecologically important EFN plant defense trait, and proposed possible adaptive hypotheses for its evolution. Creating a large database of red EFN plants, we found that red EFNs are phylogenetically, biogeographically, and morphologically widespread, but have a nonrandom distribution across both space and morphology. Using an integrative framework, we then found evidence that red pigmentation is driven by increased fungal pressure on the convergent EFN trait and suggest future work following up on these hypotheses across scales. More broadly, our study provides an example of how a set of associated convergent traits (here sugar secreting tissue and red pigmentation) may not always be the result of a single selection pressure. Rather than a unified syndrome of traits that serve the same purpose (to better attract and retain bodyguard ants), our study points to a sequential process of convergence by which an initial broader convergent phenotype – e.g. nectar to attract bodyguards to defend against herbivores – opens organisms to convergent problems – e.g. increased attack by fungal pathogens, targeted herbivory, or osmotic cellular damage – which leads to selection for a second convergent trait – e.g., red pigmentation. Ultimately, this study lays the groundwork for understanding this pattern and suggests that a red EFN may indicate a well-defended defense.

## Supporting information

Supplemental methods and results

## Acknowledgements

We thank James Latiff, Cristal Lopez Gonzalez, Allison Fletcher, and Emma Wagner for assistance in data collection; Tim Miles for providing *Botrytis* fungal cultures; Phyllis Coley, Pedro Augusto Da Pos Rodrigues, Pooja Nathan, Rosy Glos, Anselmo Nogueira, and Dale Forrister for sharing photographs of EFN tissues; Allison Fletcher for expert caterpillar identification; the iNaturalist community for recording and identifying plants with EFNs; Jill Anderson, Emily James (Georgia State Botanic Garden), University of Georgia Conservatory, and Lisa Murphy (Michigan State University Conservatory) for providing access to living specimens; Carol Kelloff and Meghann Tonner (Smithsonian herbarium), Tanisha Williams and Steven Hughes (University of Georgia herbarium), Herrick Brown (University of South Carolina herbarium), Jennifer Apland (Michigan State University herbarium), Catherine Bennett and Erin Messenger (Kew Herbarium) and Alpheus Mothapo and Erich Van Wyk (National Herbarium of South Africa) for providing access to herbarium collections; Judith Bronstein, Laural Leal, Paulo de Oliveira, and Jeff Conner for insightful discussions, Sierra Jaeger, Will Wetzel, Jeff Conner, and the Weber Lab for feedback on the manuscript, and Holly van der Stel, Kate Shaw, Brian Balcom, and Mark Hammond for crucial facilities support. This work was supported by Michigan State University Department of Plant Biology James E. Rodman endowment and Everett ‘Tex’ Beneke Fellowship, NSF GRFPs (to SME and VSP). MGW was partially supported by NSF DEB 2236747.

## Author Contributions

S.M-E. and M.G.W. conceived of the idea and designed the study. S.M-E. developed the methods; collected, analyzed, validated, and visualized the data; and wrote most of the original draft. V.S.P analyzed the phytopathogen data, wrote the corresponding methods text, and reviewed and edited the manuscript. K.J.G. and M.G.W. supervised the project and reviewed and edited the manuscript. All authors acquired funding.

## Data and Code Availability

All original data are available in the Dryad Digital Repository (DOI here after data curated). All R code is archived in Zenodo (DOI here after data curated).

