## Supplemental methods and results for "Convergent evolution of red pigmentation in extrafloral nectaries: global patterns and mechanisms"

#### EFN Location Categories

The locations consisted of “stem”, which includes all stem, tendril, bract, stipule, cataphyll, rachis, or petiole nectaries in the half of the petiole closer to the stem; “leaf base”, which includes nectaries on the base of the leaf blade (upper or lower surface), on the drip tips closest to the petiole attachment, and on the half of the petiole closer to the leaf; “upper leaf blade”, which includes all nectaries on the adaxial leaf blade, excluding those clustered at the petiole junction; “lower leaf blade”, which includes all nectaries on the abaxial leaf blade, excluding those clustered at the petiole junction; “drip tip”, which includes all nectaries on leaf tips or serrations; “leaf margin”, which includes all nectaries on entire leaf margins; and “near flower”, which includes extrafloral nectaries on the pedicel, calyx, floral bracts, phyllaries, sepals and outer surface of tepals, as well as on buds, prophylls, or developing fruit.

#### Prediction of Global Relative Phytopathogen Abundance

We trained a gradient boosting machine (GBM) (Fryda et al., 2024) package *h2o*) to predict the global relative abundance of fungal phytopathogens at the 2.5 arc minute resolution. GBM is an ensemble learning method that solves inverse-problems characterized by nonlinear relationships through sequentially fitting decision trees. In addition to the 19 bioclim variables used in Li et al. (2023), we also incorporated elevation (Fick & Hijmans, 2017), soil data (nutrient availability, nutrient retention capacity, oxygen availability to roots, excess salts, (Fischer et al., 2008), and land cover types (grass/scrub/woodland, barren/very sparsely vegetated land, urbanization, Fischer et al., 2008) as predictors. We used a 70/15/15 split (training $n$ = 9,783, validation $n$ = 2,073, testing $n$ = 2,055) and excluded observations from aquatic environments (water, sediment), human modified ecosystems (cropland, urban), and several rare habitat types (deadwood, mangrove, and lichen). Hyper-parameters were tuned based on 10-fold cross-validation and an early-stopping rule. To restrict the phytopathogen relative abundance to values between 0 and 1, we modeled it on the logit scale. The final model achieved a predictive performance of $r^2 = 0.51$ (mean absolute error = 0.060 proportion phytopathogen) on the testing dataset, indicating a reasonable ability to forecast site-level phytopathogen relative abundance and representing a substantial improvement from the $r^2 = 0.062$ reported in Li et al. (2023). We analyzed the relative abundance of the fungal phytopathogens on the logit scale (model output sale), which treats equivalent increases of phytopathogen abundance at low values as causing a greater effect on the response than at mid-ranged values. Model predictions plotted in Figure S1.

#### Fungal culture

Botrytis cinerea cultured from a commercial Michigan vineyard was grown on potato dextrose agar at room temperature.

##### *Results*

In support of the hypothesis that red EFNs contrast surrounding tissue and are thus easier to find for mutualists, we found that the distance in ant (*Myrmica*), wasp (*Polistes*), and jumping spider (*Habronattus pyrrithrix*) color space between EFNs and surrounding tissue was greater for red EFN plants than for green EFN plants (Figure S4, Table S2). This was true for both *Impatiens* (a within-species comparison between the EFN and the petiole) and for two species of *Gossypium* (comparison between the EFN and lower leaf surface in each species): *G. sturtianum* (green EFNs) and *G. australe* (red EFNs).

### Supplemental Figures and Tables

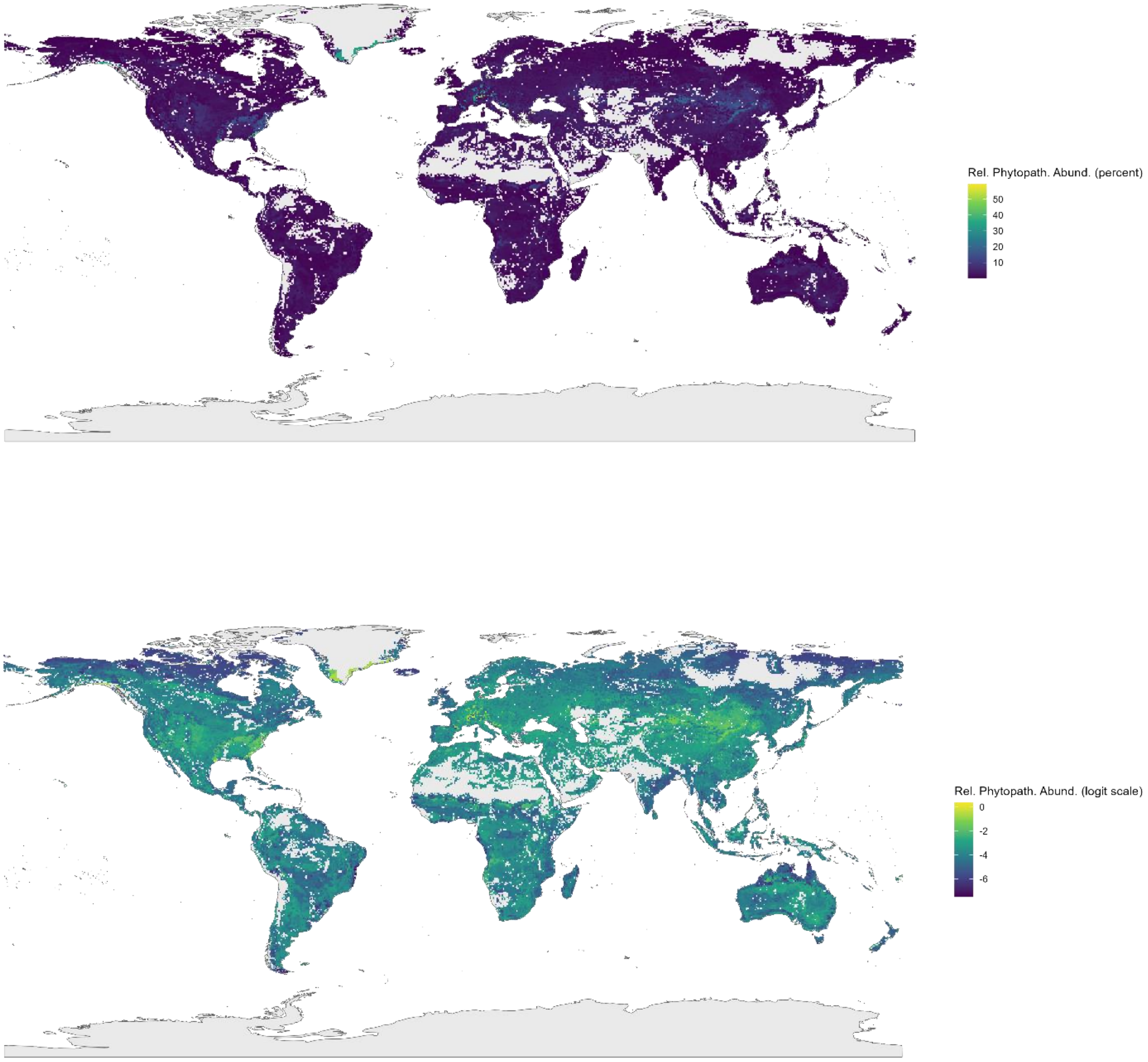


Figure S1. Model predictions of fungal phytopathogen relative abundance on the percent scale (top) and logit modeling scale (bottom). Gray areas show regions with a dissimilarity index (DI) greater than 0.4. DI per (Meyer & Pebesma, 2021)

| Family | Genus | Species | EFN Color | # in common garden | Source |
|---|---|---|---|---|---|
| Araceae | Philodendron | asplundii | 0 | - | - |
| Araceae | Philodendron | micranthum | 1 | - | - |
| Bignoniaceae | Catalpa | bignonioides | 0 | - | - |
| Bignoniaceae | Catalpa | ovata | 1 | - | - |
| Euphorbiaceae | Alchornea | glandulosa | 0 | - | - |
| Euphorbiaceae | Alchornea | trewioides | 1 | - | - |
| Euphorbiaceae | Croton | antisyphiliticus | 0 | - | - |
| Euphorbiaceae | Croton | urucurana | 1 | - | - |
| Fabaceae | Erythrina | crista-galli | 0 | - | - |
| Fabaceae | Erythrina | caffra | 1 | - | - |
| Fabaceae | Inga | pezizifera | 0 | - | - |
| Fabaceae | Inga | auristellae | 1 | - | - |
| Fabaceae | Senna | bicapsularis | 0 | - | - |
| Fabaceae | Senna | occidentalis | 1 | - | - |
| Malvaceae | Gossypium | sturtianum | 0 | - | - |
| Malvaceae | Gossypium | australe | 1 | - | - |
| Passifloraceae | **Passiflora** | **amethystina** | 0 | 30 | Kingbird Farm, Berkshire, NY |
| Passifloraceae | **Passiflora** | **caerulea** | 1 | 30 | Kingbird Farm, Berkshire, NY |
| Pedaliaceae | **Sesamum** | **indicum** | 0 | 24 | Silverhill Seeds, Cape Town, SA and GRIN Global^ |
| Pedaliaceae | **Sesamum** | **triphyllum*** | 1 | 13 | Silverhill Seeds, Cape Town, SA and GRIN Global^ |
| Polygalaceae | Monnina | richardiana | 0 | - | - |
| Polygalaceae | Monnina | exalata | 1 | - | - |
| Rosaceae | **Prunus** | **mahaleb** | 0 | 30 | Mehrabyan Nursery, Ithaca, NY |
| Rosaceae | **Prunus** | **pensylvanica**** | 1 | 5 | Fruitwood Nursery, Orleans, CA |
| Salicaceae | Populus | grandidentata | 0 | - | - |
| Salicaceae | Populus | ciliata | 1 | - | - |
| Salicaceae | Salix | pentandra | 0 | - | - |
| Salicaceae | Salix | alba | 1 | - | - |
| Viburnaceae | **Viburnum** | **dentatum** | 0 | 24 | Cold Stream Farm, Free Soil, MI |
| Viburnaceae | **Viburnum** | **rafinesqueanum** | 1 | 30 | Woody Warehouse, Lizton, IN |

Table S1. Species pairs used in herbarium infection study, with pairs also used in the common garden assays in bold. EFN color value of 1 indicates red, purple, or black EFN, while 0 indicates green or yellow EFN. In addition, the within-species pair of *Impatiens balsamina* was used in the common garden assay (30 each of red and green EFN plants, seed sourced from YEGADI Garden).

* common garden species was *S. trilobum*, due to horticultural availability.
** common garden species was *P. avium* due to horticultural availability.
^ GRIN accessions in data files archived in Dryad

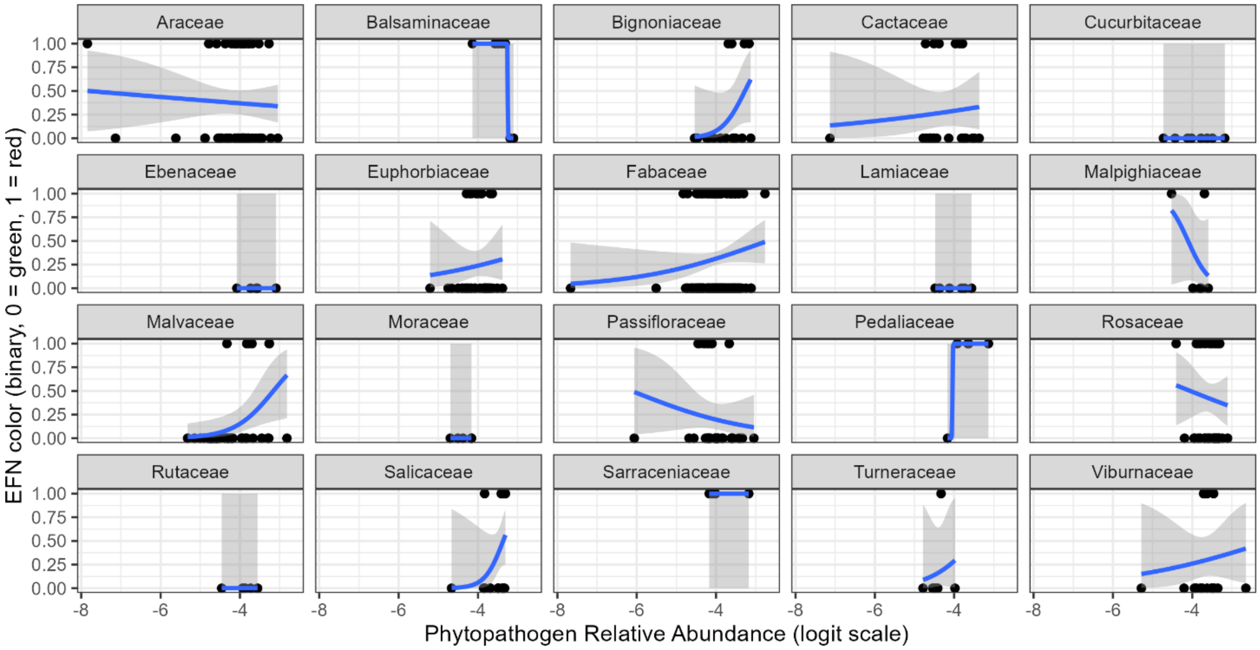


Figure S2. Median predicted phytopathogen relative abundance across the species range predicting EFN color, faceted by family (includes only families with 5 or more species in the dataset).

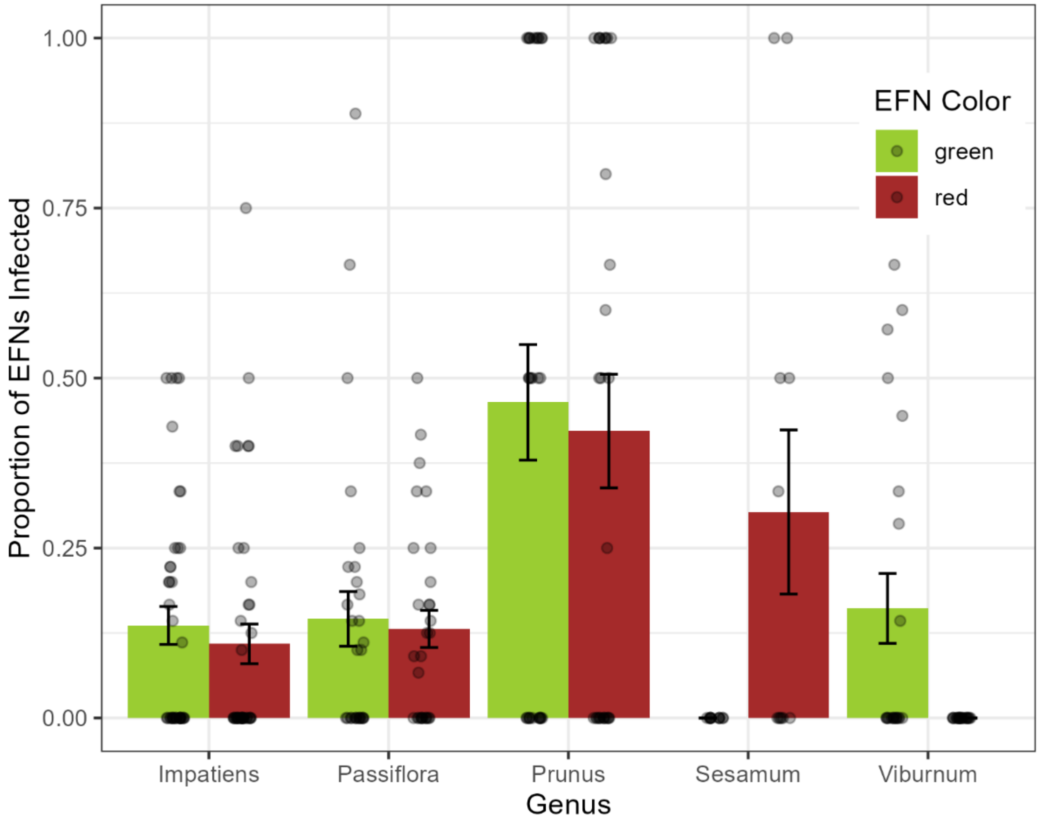


Figure S3. Across species pairs, we observed an overall higher rate of incidental infection of EFNs on leaves fed to crickets in humidified petri dishes. The green EFN species bar is shown on the left of each pair. Note that due to limited survival of *Prunus avium* (red EFNs), the majority of leaves of this species used in the cricket bioassay are from mature trees growing around Kellogg Biological Station.

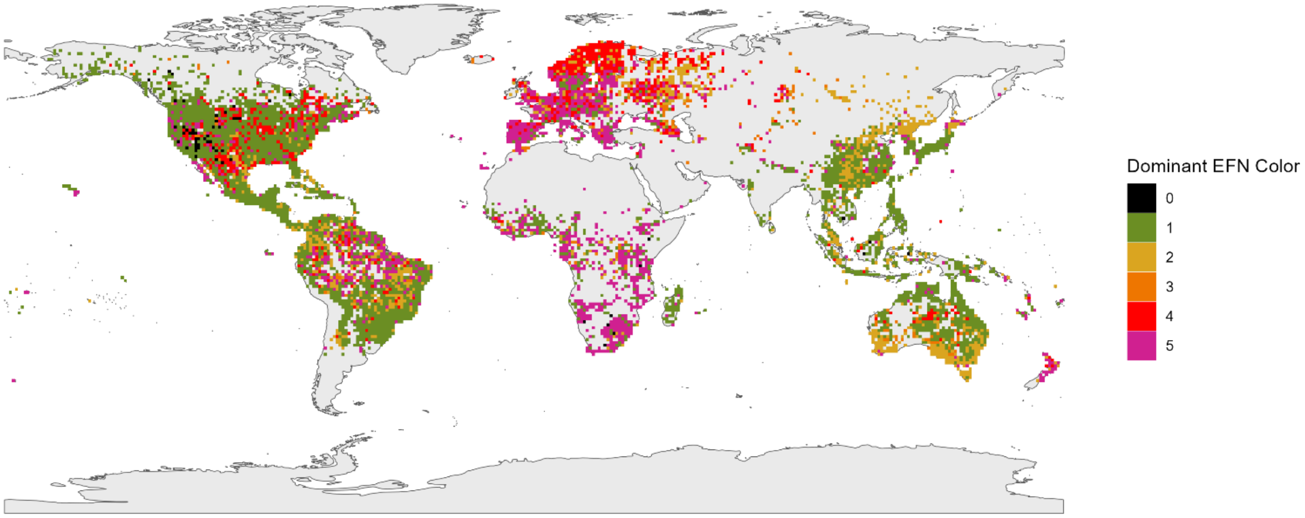


Figure S4. Map of dominant EFN color within each 1 degree raster cell across the GBIF dataset (data subsampled to one occurrence per species per cell). 0 = brown EFNs, 1 = green, 2 = yellow, 3 = orange, 4 = scarlet, 5 = maroon. Multiple EFN color scores within each species (e.g. species with more than one EFN type, or with EFN that ranged from yellow to red) collapsed to single maximum score. Figure is faceted below for accessibility.

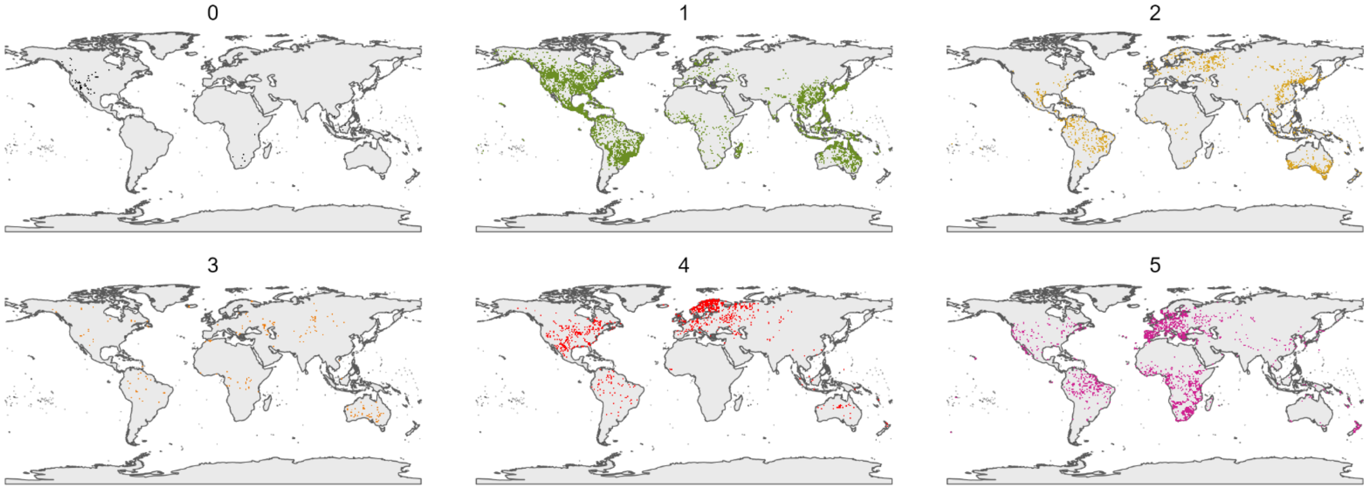
